# Cultural determinants of the gap between self-estimated navigation ability and wayfinding performance: evidence from 46 countries

**DOI:** 10.1101/2022.10.19.512889

**Authors:** S. Walkowiak, A. Coutrot, M. Hegarty, P. F. Velasco, J. M. Wiener, R.C. Dalton, C. Hölscher, M. Hornberger, H. J. Spiers, E. Manley

**Author notes:** These authors contributed equally to this work (H. J. Spiers, E. Manley).

## Abstract

Cognitive abilities can vary widely. Some people excel in certain skills, others struggle. However, not all those who describe themselves as gifted are. One possible influence on self-estimates is the surrounding culture. Some cultures may amplify self-assurance and others cultivate humility. Past research has shown that people in different countries can be grouped into a set of consistent cultural clusters with similar values and tendencies, such as attitudes to masculinity or individualism. Here we explored whether such cultural dimensions might relate to the extent to which populations in 46 countries overestimate or underestimate their cognitive abilities in the domain of spatial navigation. Using the Sea Hero Quest navigation test and a large sample (N=383,187) we found cultural clusters of countries tend to be similar in how they self-rate ability relative to their actual performance. Across the world population sampled, higher self-ratings were associated with better performance. However, at the national level, higher self-ratings as a nation were not associated with better performance as a nation. Germanic and Near East countries were found to be most overconfident in their abilities and Nordic countries to be most under-confident in their abilities. Gender stereotypes may play a role in mediating this pattern, with larger national positive attitudes to male stereotyped roles (Hofstede’s masculinity dimension) associated with a greater overconfidence in performance at the national level. We also replicate, with higher precision than prior studies, evidence that older men tend to overestimate their navigation skill more than other groups. These findings give insight into how culture and demographics may impact self-estimates of our abilities.

## Introduction

It is important to be able to estimate one’s own cognitive abilities accurately. If we underestimate our capacity for a task, we may preclude ourselves from beneficial outcomes. On the other hand, overestimating our abilities can be dangerous if we are then unable to cope with the required challenges. One such challenge all humans face is spatial navigation. There is evidence that self-reported navigation ability is related to objective measures of this ability [1–4]. A number of studies have shown this relationship using standardised measures of self-reported spatial abilities, such as Santa Barbara Sense of Direction Scale [5] or Wayfinding Self-Efficacy Questionnaire [6]. The relation holds both on an individual level and on a group level: men tend to perform better on navigation tasks [7–10], and, as a group, also to self-rate themselves as better navigators [11]. Beyond the topic of navigation, previous studies have found that positive self-estimates of ability are associated with better outcomes in tasks such as motor learning [12, 13], game-based learning environments [14], science assessments [15], and overall academic success [16]. These self-estimates are often viewed from the standpoint of self-efficacy: how estimates go on to predict outcomes [17].

While self-estimates are related to navigation ability, the relationship between the two is complicated by additional evidence that some groups of people may over- or under-estimate their navigation performance. These findings do not deny the well-established relation between self-reports and navigation ability. Rather, what they indicate is that there must be other factors that influence self-reports of navigation ability besides navigation ability itself. Given the dynamic and multi-dimensional nature of the relationship between self-reports and navigation ability, it is likely that between-subject factors (e.g. gender, education, age) and cross-cultural differences (e.g. gender stereotypes, country-specific education, social policies, etc) affect both self-estimates and performance in wayfinding tasks [18, 19].

Cross-cultural effects are likely to play a role in the efficacy of self-estimates. For instance, in some cultures, it may be seen as a positive to present oneself as competent at tasks, while in others it may be important to be self-effacing [8, 19, 20–28]. Existing research also indicates that gender and age factors are important elements influencing the reliability of self-estimates. Previous studies have reported that older adults and men tend to overestimate their navigation abilities [22, 29–31]. This overestimation reported by older adults is often associated with a possible lack of awareness of their cognitive decline, which becomes especially noticeable in laboratory-based settings, navigation tasks implemented in unfamiliar places or using modern virtual reality (VR) software [32–34]. Other studies have found that females tend to rate their spatial skills as lower compared to men even when there is no objective difference in performance [30, 35–39]. To explain this gender disparity, Cross et al. [36] attributed female lower spatial self-evaluation to women’s higher susceptibility to social influences and their greater dependency on social information. As spatial skills have been traditionally regarded as more masculine qualities [40], females are more likely to conform to this gender-stereotyped expectation of their own abilities and consequently, they tend to respond more modestly to self-estimate questions in spatial, navigating or wayfinding tasks. The awareness of these negative stereotypes of women’s abilities may in turn influence female performance on these tasks. Nori & Piccardi [35] reported however that in some cases in which the overall population of individuals claims high levels of self-efficacy on a typically masculine task, gender differences disappear if women share the general belief of high competency on the specific task (e.g. in studies with Italian Air Force pilots, [41, 42]). This positive self-perception may be caused by more assumed ‘masculine cognitive characteristics’ related to analytical, rationale, and mathematical thinking [43] of women in such populations, which appears to be predictive of their better overall wayfinding ability [44].

We hypothesise that the impact of cultural norms on self-estimates in women also applies to larger cross-national samples where self-estimates are likely to be affected by country-specific stereotypic gender beliefs, societal norms and cultural variation. In support of this claim, Oettingen [45] argued that common value systems to which individuals living in different countries or geographical regions are exposed during their lifetime can contribute to their self-evaluation on a particular task. When characterising the role of these correlates, she pointed to six cultural dimensions (power distance, individualism, masculinity, uncertainty avoidance, long-term orientation and indulgence) formulated by Geert Hofstede [46, 47] as the theoretical framework for understanding between- and within-countries variations in self-reported abilities and their effects on task performance. Oettingen’s theory has been applied in studies explaining the effects of selected cultural determinants of self-beliefs such as individualism vs. collectivism dimension in various fields e.g. education and business [48, 49], and recently cultural factors were found to be a significant predictor of the academic attributional style in a study of British and Turkish students [50], maths self-evaluation, anxiety and performance on maths tests across 41 countries [51], maths competence across 34 countries [52], intelligence and differences in self-estimates between US and Nicaraguan children and adolescents [53], as well as academic achievement, learning styles, and teacher and student self-beliefs [54, 55]. Most of the referenced studies agree that self-estimates are predictive of performance across different countries but that cultural dimensions and various other socio-demographic factors may additionally influence how self-evaluation influences individual or group behaviours on different types of tasks.

Cross-cultural and between-countries analyses in the field of human navigation are rather scarce. The ones that exist often suffer from small sample sizes and limited geographical coverage. And while there are some studies that investigate general spatial abilities and cognition, these do not include route and wayfinding tasks. Because of the limitations in existing studies, the question remains of how much of the differences in self-reports of navigation ability are due to actual ability and how much they are due to other factors such as age, gender and cross-cultural differences. This question is difficult to answer because it is challenging to test many people across diverse cultures using traditional methods.

Here, we overcome this challenge by testing over 4.3 million individuals with our navigation task embedded in the videogame app Sea Hero Quest. Using Sea Hero Quest, we are able to explore the impact of quantified cultural dimensions on the gap between wayfinding performance and self-estimated navigation ability across 46 countries.

## Results

Wayfinding performance was measured using the Sea Hero Quest mobile app [19]. This test of navigation is embedded in a video game in which the players steer a virtual boat through nautical environments seeking mystical sea creatures (see Figure 1). Position was tracked during navigation and the performance calculated from the distance travelled (see Methods: Metrics and Statistical Analysis section for details). Self-estimates were collected along with other demographics in the app via a question that asked if players judged their ability to navigate was: very good, good, bad, or very bad.

**Figure 1.**
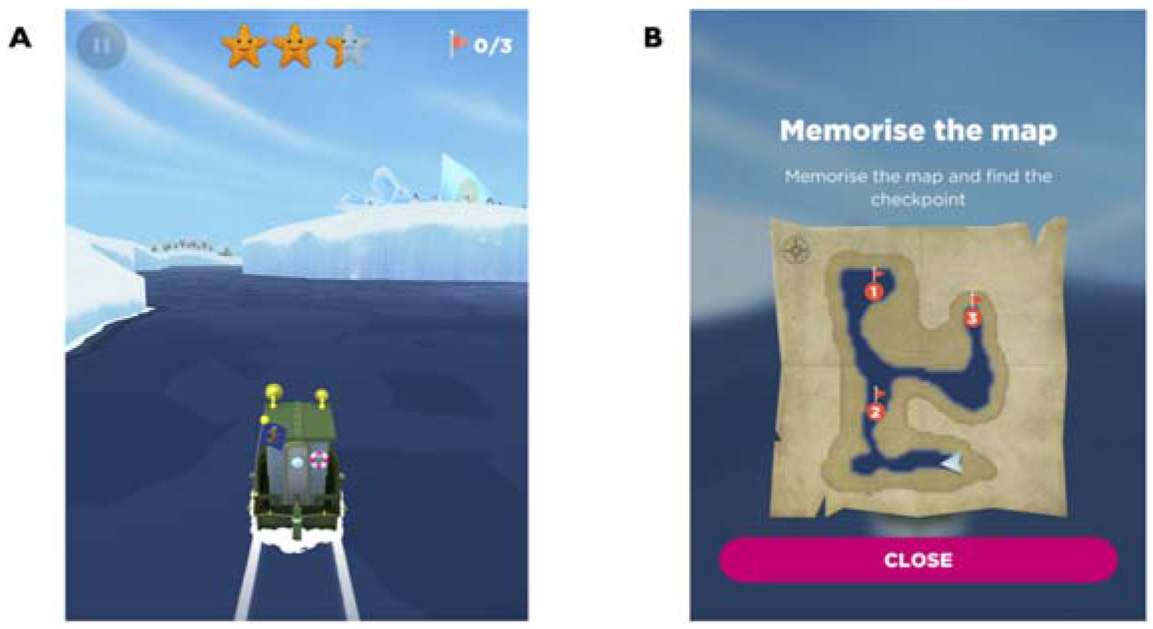
Navigation task Sea Hero Quest. A) View during active navigation (level 11). B) Map viewed before active navigation of level 11, indicating start position (triangle) and checkpoints (numbered circles).

### Age, gender, self-estimates and wayfinding performance

Following the data processing and sampling operations described in the Methods section, the resulting sample size used to explore the relationships between age, gender, self-estimates of navigating skills and wayfinding performance included 383187 individuals (from 46 countries) who completed six wayfinding levels of the Sea Hero Quest game: 3, 6, 7, 8, 11 and 12 (*N_Females_* = 172969, *MeanAge_Females_* = 37.08 ± 14.53, *N_Males_* = 210218, *MeanAge_Males_* = 37.14 ± 13.39; for more details, see Supplementary Information, Appendix A, Table S1).

The ordinal logistic regression analysis (with self-estimates nominal responses as the dependent variable; gender and age bands as the independent variables) revealed that males were significantly more likely to report good navigating skills than females (*LR* = 21895.2, *p* < .001). Based on the model (Supplementary Information, Appendix A, Table S2), men were almost two times more likely than females to rate themselves as very good navigators and approximately half as likely as women to rate their navigating skills as either very bad or bad across all age bands (Figure 2A). Age was also found to be a statistically significant positive predictor of navigating self-estimates (*LR* = 1299.5, *p* < .001). The best self-reported navigation ability in the sample was reported by 40-59-year-old males (Figure 2B), however older males (between 60 and 70 years old) rated their navigating skills more favourably than the youngest men in the study (19-29-year-olds) despite their much poorer actual wayfinding performance when compared to all younger age bands of the male sample (Figure 2C). This finding supports the results of earlier studies [22, 29] which found similar overestimation of navigating abilities amongst oldest male participants. With our larger sample we can now reveal that at each age band there is a preserved ordering of self-ratings in relation to performance, e.g. people who self-rate as very good are the best navigators (Figure 2C). This provides evidence that when people self-rate they may do so in relation to their peers, both with regard to people of the same age and people of the same gender.

**Figure 2.**
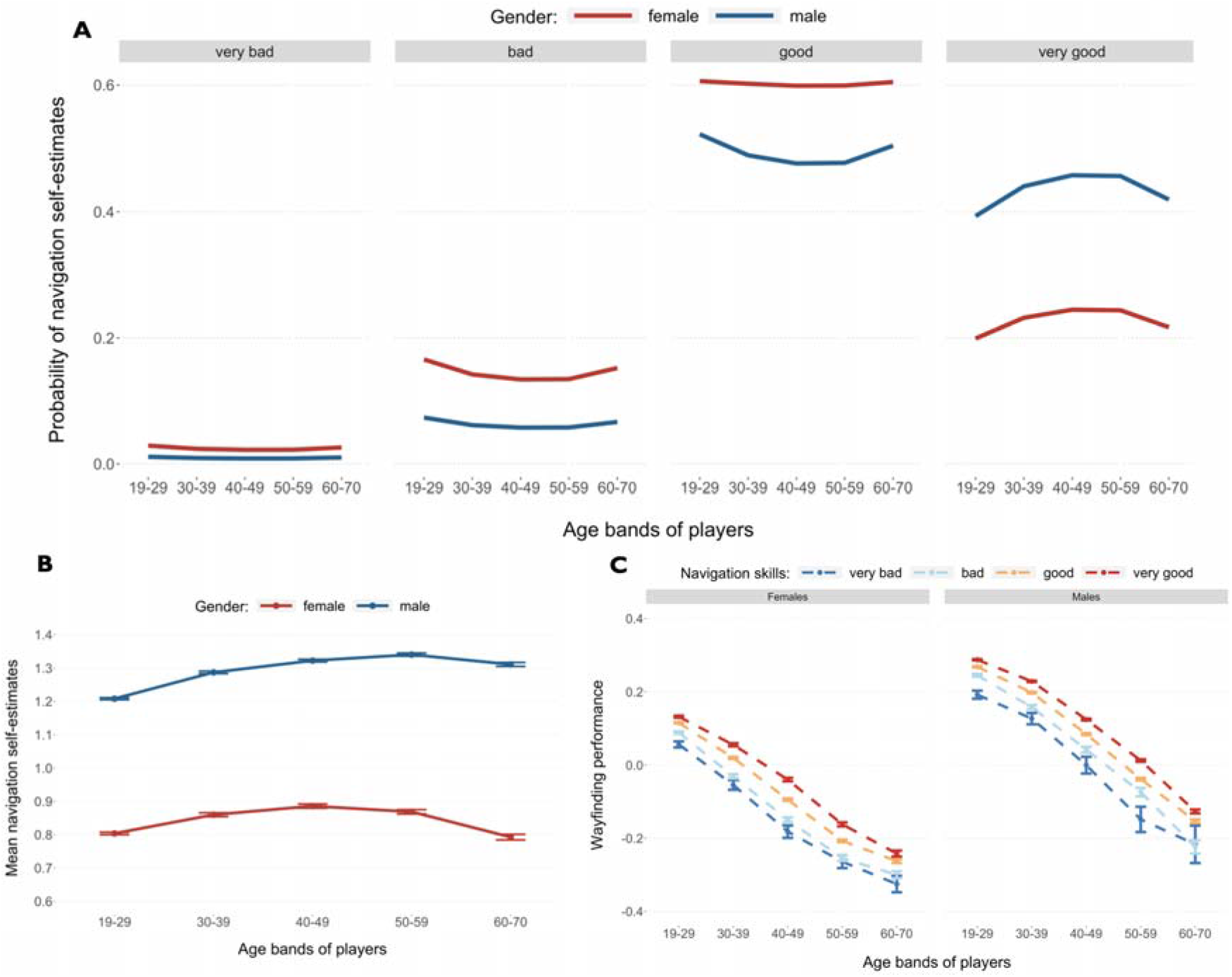
Self-estimated navigation skills and wayfinding performance by gender and age. A) Probabilities of reporting specific navigation self-estimates by females and males across five age bands represented in the sample. B) Average self-estimates of navigation skills (with standard error bars) for males and females across five age bands. C) Wayfinding performance (with standard error bars) of both female and male players for six SHQ wayfinding tasks (Levels 3, 6, 7, 8, 11, and 12) across five age bands.

We used a hierarchical multivariate linear regression analysis (Supplementary Information, Appendix A, Tables S3 and S4) to establish whether the reported ratings of navigation skills could predict wayfinding performance as the dependent variable. A model with self-ratings included captures more of the variance than a model without them (*F*_3_ = 536.73, p < .001) with self-rated ability the fourth strongest predictor of wayfinding performance amongst all independent variables used in the model (*F*_3, 370561_ = 536.73, *p* < .001, *η*^2^ = 0.004), behind age (*F*_1, 370561_ = 55125.38, *p* < .001, 2 = 0.123), gender (*F*_1, 370561_ = 19685.08, *p* < .001, *η*^2^ = 0.044) and home environment (*F*_2, 370561_ = 587.35, *p* < .001, *η*^2^ = 0.003), but before education (*F*_3, 370561_ = 361.11, *p* < .001, *η*^2^ = 0.002) and commute time (*F*_2, 370561_ = 61.27, *p* < .001, *η*^2^ = 0.0003).

### Cultural determinants of self-estimates and wayfinding performance

To explore how cultural differences might relate to self-ratings, we aggregated the data from the 46 countries sampled into the 11 cultural clusters derived by Ronen & Shenkar [56] (Figure 3A; also see Methods for details): Germanic, Eastern European, Nordic, Far East, Near East, Confucian Asia, Latin Europe, Eastern Europe, Anglo, Arabic and African. The highest proportions of population with either good or very good self-estimates of their navigating skills were recorded in Germanic, Eastern European and Latin American cultural clusters (96%, 90% and 88% of their respective samples), whereas groups of countries with the most modest beliefs about their navigating skills were Latin Europe, Nordic and Confucian Asia (85%, 84%, and 81%, respectively; proportions of self-reported navigation skills by country and cultural cluster available in the Supplementary Information, Appendix A, Tables S5 and S6). However, more positive self-estimates did not translate to better wayfinding performance at the national level; there was no significant correlation between these two metrics (Figure 3H). Notably, we replicated our prior finding that GDP correlates with navigation performance [19] (here focusing on wayfinding in 46 countries) Spearman’s *rho*=0.572, p<.001. By contrast self-estimates do not correlate with GDP (Spearman’s *rho*=0.11, p=.48). Thus, it appears that increased economic wealth in a country is associated with better performance, but not better self-estimates of performance.

**Figure 3.**
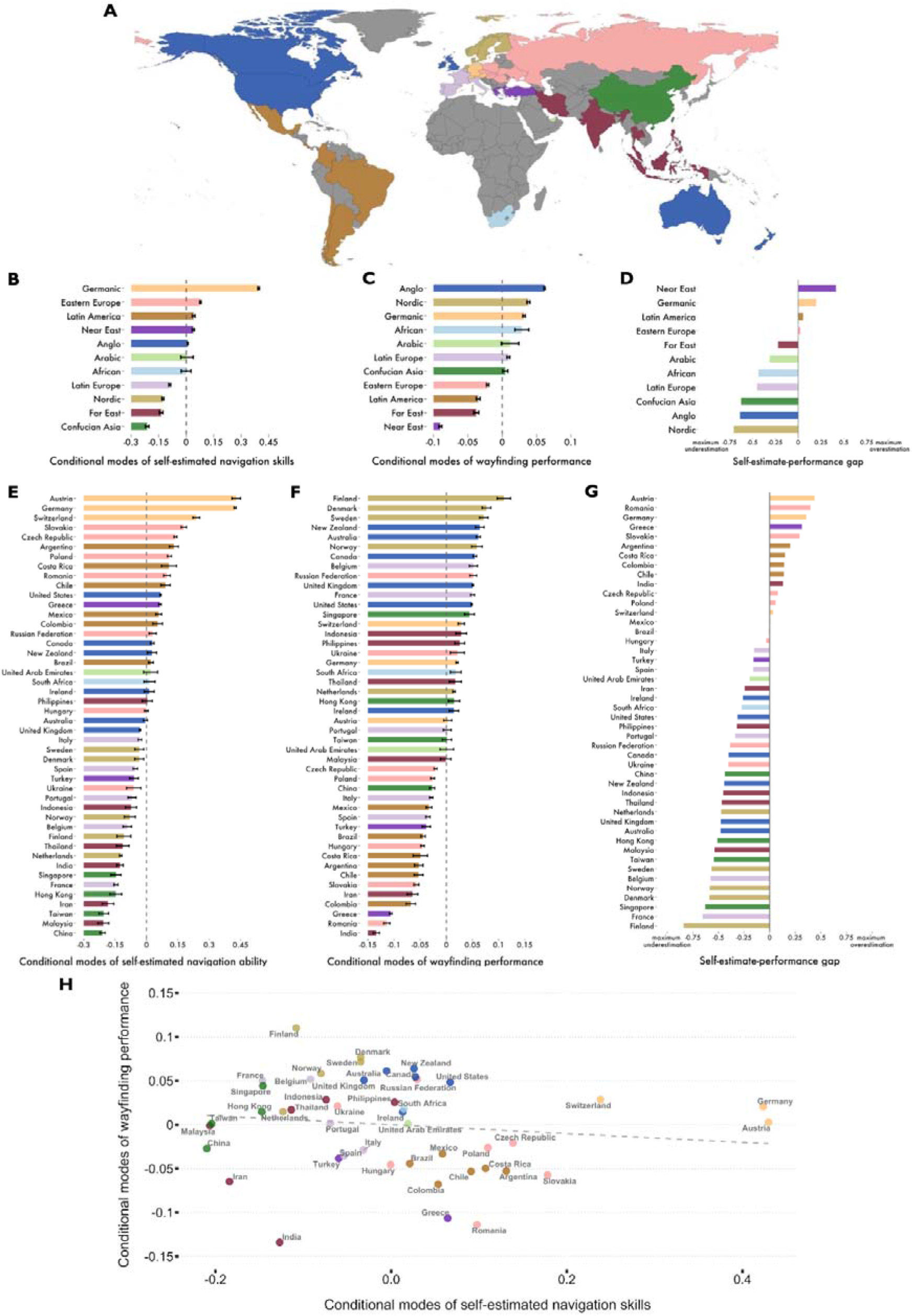
Self-estimated navigation skills vs. wayfinding performance across countries and cultural clusters. A) Global map of cultural clusters as defined by Ronen & Shenkar [56]. Conditional modes (controlled for age and gender) of self-estimated navigation skills for cultural clusters (B) and countries (E) with indicated standard errors. Conditional modes (controlled for age and gender) of measured wayfinding performance for cultural clusters (C) and countries (F) with indicated standard error bars. Self-estimated ability vs. wayfinding performance gap by cultural cluster (D) and country (G) estimated by calculating the difference between min-max normalised conditional modes of navigation skills and wayfinding performance (range [−1,1] where −1 denotes maximum possible underestimation, whereas 1 indicates maximum overestimation). H) National self-estimated navigation ability and the wayfinding performance are not significantly correlated (Spearman’s *rho*=−0.2, p=.181). See Supplementary Information, Appendix A, Figure S1 for results split by gender.

We next explored the gap between a country’s self-rated performance and actual performance. We found that similar to country clusters of self-ratings, country clusters also showed tendencies to be over or under confident as a group. Near Eastern and Germanic countries were most likely to overestimate their ability and Nordic countries most likely to underestimate their ability (Figure 3). Examining the geographic spread of the data further highlights the tendency for neighbouring countries and countries within cultural groups to be more similar (Figure 4). We found a similar pattern for both men and women when the linear models were fit to men and women separately (Supplementary Information, Appendix A, Figure S1). Thus, overestimation of performance in some countries is not due to, for example, predominantly the men overestimating their performance and female participants accurately estimating them.

**Figure 4.**
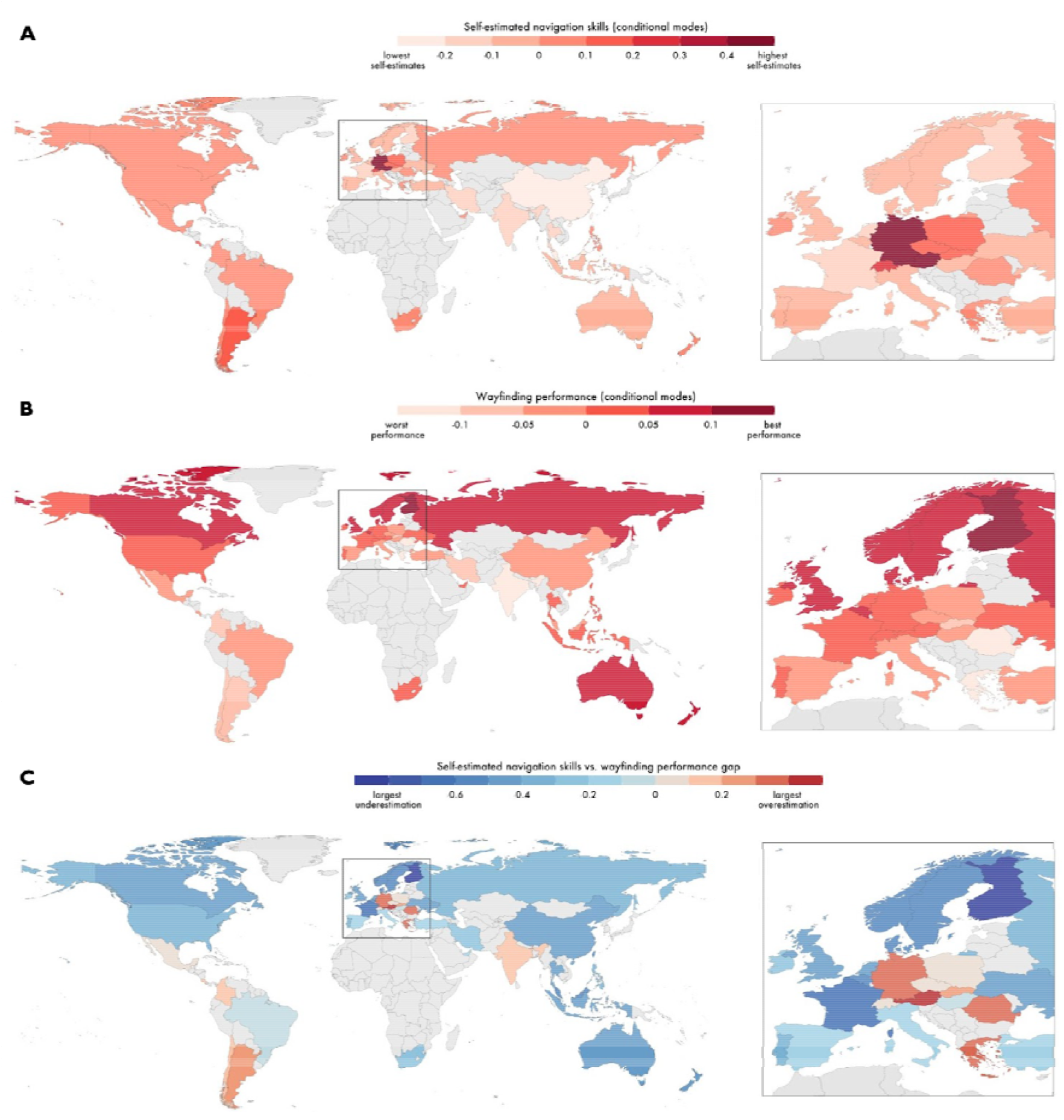
Global representation of self-estimated navigation ability, measured wayfinding performance and the self-estimate-performance gap. A) Self-estimated navigation skills by country; players from countries in darker red colour reported higher self-estimates. B) Wayfinding performance measured during the navigation tasks; countries filled with darker red colour performed better. C) Self-estimated navigation skills vs. measured wayfinding performance gap; players from countries filled with light pink to dark red colours overestimated their navigation skills, whereas countries in darker blue largely underestimated their navigation ability. Countries with no data filled with light grey.

Based on past literature [43, 44], we considered that one reason some countries might overestimate ability is that they have a stronger association with male typical stereotypes, with navigation one of these stereotypes. If a country feels it is important to be good at male associated activities it is possible that nationals of that country would be more confident in these abilities. In support of this we found that greater masculinity ratings for a country was associated with greater overconfidence of its population (Spearman’s *rho*=0.43, p=.004; Figure 5). We also found a similar association with purely self-estimate ratings (Spearman’s *rho*=0.40, p=.008). To explore more broadly the cultural dimensions that might impact the gap we examine the full set of 6 metrics developed by Hofstede [46, 47] (Supplemental Information, Appendix B, Table S7). We found that masculinity was the strongest predictor of the gap and survived Bonferroni correction for significance in this model. We also found that Hofstede’s cultural dimension of uncertainty avoidance was also a predictor of the gap in this statistical threshold corrected model. Nations who are more keen to avoid uncertain situations were more likely to overestimate navigation ability (see Supplemental Information, Appendix B).

**Figure 5.**
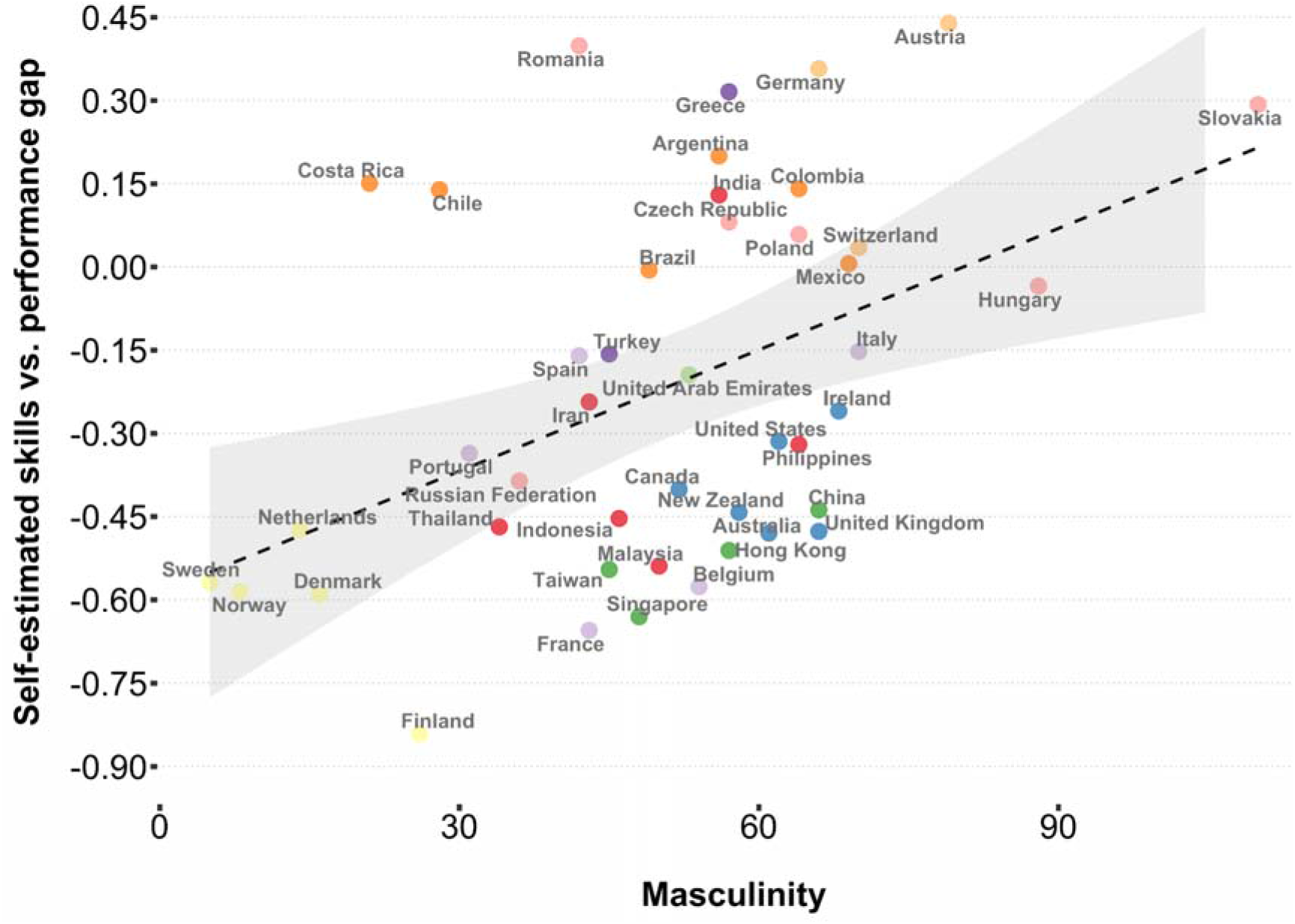
Relationship between the self-estimate-performance gap and Hofstede’s masculinity metric (Spearman’s *rho*=0.43, p=.004).

Because we found differences between men and women in self-ratings we explored how national patterns differed for men and women. We found relatively consistent patterns for both men and women in countries, for countries with high self-ratings, both women and men rated themselves as high, and for countries with low self-ratings, similar low self-ratings were present for men and women. Thus, the patterns observed at the national level do not appear to be strongly influenced by gender groups, but rather than by overall cultural attitudes.

Having previously reported that the navigation performance gap between men and women was correlated with the gender gap index [19], we also explored the self-ratings - performance gap in relation to such a metric. We found there was significant positive correlation between the self-rating / performance gap and the gender inequality index of a country (Spearman’s *rho* = 0.40, *p* = .007). Thus, the more unequal a country in terms of gender gap, the greater the overconfidence.

## Discussion

Here we explored variation across nations in terms of their population’s self-estimates at navigating, their ability to perform a virtual navigation task and the gap between these two measures. Extending in more depth than prior studies, we found that older men were more likely to overestimate their ability. We now report that, while across the global sample, performance across all participants was related to pooled self-estimates of navigation ability, at the national level a nation’s self-estimates and performance were not correlated. We found that participants in Germanic countries were much more likely to report high self-estimates than other countries, whereas Confucian Asian countries reported the lowest self-estimates. Near Eastern countries showed the largest over-estimation in their abilities, whereas Nordic countries showed the greatest under-estimation in their abilities. We found that across nations the gap between self-estimates and performance was associated with the extent of affirmation for gender roles, the greater the association with masculinity in a country, the more likely citizens of that country were to overestimate performance. We discuss how these findings advance understanding of how self-estimates of ability relate to actual performance for a core human ability: spatial navigation.

Consistent with prior research, we found that across the entire sample male participants were twice as likely as females to rate themselves as very good navigators [5, 35–39]. Men also displayed a much lower tendency to report their navigating skills as either bad or very bad. Middle-aged and older participants generally held more favourable views about their navigating abilities than younger adults and this effect was particularly strong in the male group. For example, the oldest males in the sample (*i.e.* 60-70-year-olds) rated their wayfinding skills higher than the youngest 19-29-year-old men despite their performance being much worse than all younger males including the youngest age group. This finding is congruent with the research from van der Ham et al. [30, 31] who reported that overestimation of spatial navigation abilities increases with age, with this more common for males than females. It also validates the earlier research by Taillade et al. [22, 29] who found that older individuals rated their spatial abilities more positively than younger individuals, but they also displayed poor accuracy in their self-estimates due to age-related decline in performance.

We also replicate our past finding that the GDP of countries correlates with the performance of these countries [19], and now extend this to reveal that GDP is not correlated with self-ratings. Thus, while the economic wealth of a country (raising education, health care and access to travel) appears to enhance cognition, it does not relate to the way in which a population will self-reflect on their abilities.

Our results provide the largest cross-cultural study of the difference between self-reported navigation skill and ability. While across the whole world population higher self-estimates are associated with better performance, we found no correlation between a nation’s average performance and a nation’s self-estimated ability. Germanic countries were found to self-rate their abilities much more positively than other countries, with other cultural clusters (such as Nordic nations) showing similar overall ratings scores for self-estimates. If self-ratings for navigation are indicative for self-ratings in general, it would suggest Germanic cultures value taking a positive attitude to self-achievement at tasks, while Nordic nations value modesty in considering one’s abilities. We found striking examples of underestimation were countries from the Confucian Asia and Far East, which despite reporting very modest ratings of their navigating abilities performed well in comparison to other cultural clusters. On the other hand, countries representing the Germanic cluster were more susceptible to overestimation: they performed relatively poorly while expressing very positive beliefs about their spatial navigation abilities. We hypothesised that, due to past evidence for gender bias in self-estimates studies of navigation [41, 42], that the attitude towards *masculinity* in the population might manifest in bias in the self-estimates - with high masculinity associated with higher self-estimates. We found that both the population self-estimates and the over/under-estimation in performance were associated with population attitudes to masculinity. This finding supports Oettingen’s [45] theory that a domain-specific sense of competence, in this case wayfinding, is likely to vary between countries and cultures based on the shared value systems which an individual has experienced. We show here this extends not only to a self-reported estimate, but to over- and under-estimation of performance. Notably, we found little evidence that such relationships with masculinity were different for men and women. Thus, generally, it is not the self-estimates of men or the self-estimates of women within a country (or their gap to performance) that underlie the association with masculinity, but collective populations.

Exploring beyond masculinity as a metric we examined other metrics collected by Hofstede [46, 47] in a model with corrected thresholds and found that *uncertainty avoidance* was also significantly correlated with overestimation of abilities, albeit less so than masculinity. This indicates that people in countries where certainty is valued have a tendency to consider themselves good navigators, whether or not they, as a group, are. It is currently unclear why this might be and future research will be useful to explore this.

A limitation to the current research is the treatment of culture at the nation-state level. Within most nations, different cultures exist, and for many nations, different languages are spoken by sub-populations. Prior research has shown that language, attitudes to childhood, and the environment occupied by sub-groups of a nation can have an impact on spatial cognition [18, 61–65]. Thus, it seems likely that both the language and the environment of relevant sub-groups will affect the relationship between self-ratings and navigation ability at a population level. One further challenge is that different cultural groups will differ in access to technology and smart-phones, and our current results are limited to a self-select group of participants who have access to such technology. However, our current results are unlikely to be mediated by such effects since Germanic and Nordic countries both have good access to technology and occupy different ends of the self-rating scale for nations. Future research involving more traditional cultures will require careful consideration regarding the use of technology in the assessment of cognition.

In summary, here we reveal world-wide patterns in self-ratings and over/under-confidence of a core cognitive skill: navigating. We replicate prior age and gender patterns with significantly more power than previous studies. We find that attitudes to masculinity across a nation are associated with self-estimates in ability and their over/under-estimation. Future research will be useful to extend these findings to other cognitive domains.

## Methods

### Data and subject details

#### Sea Hero Quest game design

The research questions were tested using a large-scale dataset collected via Sea Hero Quest – a mobile game application designed to obtain benchmarks of typical spatial abilities and navigation behaviours of healthy populations worldwide. The game contains eighty wayfinding, path integration, radial maze and other, research-unrelated, game engagement levels along with additional optional socio-demographic questions which asked players about their age, gender, country, education, rating of their navigation skills, hand used for writing, the environment they grew up in as well as their average daily travel time and a typical amount of sleep at night. A detailed description of the game design is provided in Coutrot et al. [19]. The ecological validity of Sea Hero Quest has already been confirmed by Coutrot et al. [57] who found that the navigational performance recorded using the game reflects real-life wayfinding behaviour.

The full Sea Hero Quest dataset contains behavioural and socio-demographic data collected between April 2016 and April 2019 from over 4.3 million players globally with over 50 million gameplays recorded across all wayfinding and path integration tasks. Access to the full dataset is described in the Data Availability statement of this manuscript.

The starting screen of the Sea Hero Quest game included debriefing and consent sections with clearly identified goals of the study as well as the purpose and extent of data collection. Additionally, the full explanation of the study was provided within the gameplay (via the ‘journal’ icon) and the consenting players could access this section and withdraw from playing at any point of the game.

#### Participants and sample size

A subset of 383187 individuals (representing 46 countries; *N_Females_* = 172969, *MeanAge_Females_* = 37.08 ± 14.53, *N_Males_* = 210218, *MeanAge_Males_* = 37.14 ± 13.39) out of the total number of over 4.3 million people who downloaded and played the Sea Hero Quest mobile game application was used in the analysis. The final sample was composed of individuals from 19 to 70 years old who completed six wayfinding levels of the Sea Hero Quest game: 3, 6, 7, 8, 11, and 12, provided their socio-demographic information and responded to the question related to their self-estimated navigation skills. Additionally, the sample was restricted to include only these participants who represented countries with at least 500 qualifying players and excluded the outliers. Full sampling methodology is described in the *Data processing, sampling and controlling for fake demographics* of Methods. The country-by-country descriptives of the sample are provided in the Supplementary Information, Appendix A, Table S1.

#### Wayfinding tasks

For the purpose of this study, we only used a selection of wayfinding levels in which the overall goal was to navigate a boat from the origin to the destination in various spatial layouts. They included a number of checkpoints (represented as buoys) which players had to visit in a specific sequence in order to complete the task. Before each game level, players were first shown a map of the corresponding layout environment which they could look at for as long as they needed. At some levels the maps were obscured which forced players to learn the layouts by exploration and made the navigation tasks more difficult. The first two levels of the game were designed as tutorials to allow players to familiarise themselves with the game controls. The performance on these two tasks was used in the analysis to account for any variation in computer experience amongst the players.

#### Cultural clustering of countries

One of the focal points of the analysis was to assess the differences in wayfinding performance and navigating self-estimates between countries as well as groups of countries clustered together based on their cultural similarities. In order to determine the groups of culturally similar countries we followed the cultural clustering implemented in the GLOBE 2004 study [58] with further extensions proposed by Ronen & Shenkar [56]. The original GLOBE 2004 project grouped 52 still existent countries into 10 culturally distinct clusters based on cultural dimensions such as power distance, uncertainty avoidance, societal institutional collectivism, assertiveness, gender egalitarianism etc. which vastly overlapped previously introduced Hofstede’s cultural dimensions [46]. Ronen & Shenkar [56] additionally analysed clusters defined in 11 most influential studies (including Hofstede’s study and the GLOBE project) and proposed 11 global and 15 consensus clusters to group 70 distinct countries. In this study, the Ronen & Shenkar’s solution has been employed to group 46 countries represented in the sample into 11 global clusters as displayed in Table 1. As some countries (*i.e.* Albania, Croatia, Egypt, Lebanon, Macedonia, Puerto Rico, Saudi Arabia, Serbia, and Vietnam) never featured in the GLOBE or Ronen & Shenkar’s studies, their cluster membership was problematic and therefore we have removed players who represented these countries from further analysis.

**Table 1.** Ronen & Shenkar’s cultural clusters for 46 countries represented in the sample.

| **Cluster name** | **Countries** |
| --- | --- |
| African | South Africa |
| Anglo | Australia, Canada, Ireland, New Zealand, United Kingdom, United States of America |
| Arabic | United Arab Emirates |
| Confucian Asia | China, Hong Kong, Singapore, Taiwan |
| East Europe | Czech Republic, Hungary, Poland, Romania, Russian Federation, Slovakia, Ukraine |
| Far East | India, Indonesia, Iran, Malaysia, Philippines, Thailand |
| Germanic | Austria, Germany, Switzerland |
| Latin America | Argentina, Brazil, Chile, Colombia, Costa Rica, Mexico |
| Latin Europe | Belgium, France, Italy, Portugal, Spain |
| Near East | Greece, Turkey |
| Nordic | Denmark, Finland, Netherlands, Norway, Sweden |

### Metrics and statistical analysis

#### Measuring self-estimates of navigation skills

The self-estimates of one’s navigation skills were collected from the players via the optional question *“How good are you at navigating?”* asked after completion of the second wayfinding task of the game (Level 2). The available answers were distributed on a four-item Likert-type scale without a neutral/average score: *‘very bad’, ‘bad, ‘good’*, and *‘very good’*. The obtained responses were used as a proxy for measuring the self-reported navigation ability. Our analysis treated the self-estimates responses as either categorical (as recorded by the game) or numeric values. A numeric version of this variable was obtained by assigning a numeric value to each response according to the following coding: *‘very bad’* = −2, *‘bad’* = −1, *‘good’* = 1, *‘very good’* = 2. This numeric-type measurement of navigation skills was applied to quantify the average (arithmetic mean) self-estimates for each gender, age band and country as well as to calculate the country-specific gap between the self-reported ability and measured wayfinding performance.

The average self-estimated navigation skills for each country and cultural cluster in the sample were estimated with conditional modes by extracting the intercepts for 46 countries in the sample and 11 cultural clusters from linear mixed models with fixed effects for age and gender and random effect for country (or cluster, respectively): Self-Estimated Navigation Skills ~ Age + Gender + (1 / Country) and Self-Estimated Navigation Skills ~ Age + Gender + (1 / Cluster). We used lme4 [59] and lmerTest [60] packages for the R programming language to implement these models.

#### Wayfinding performance metrics

Measuring wayfinding performance of individual gameplays followed the methodology of correcting distance presented by Coutrot et al. [19] and was estimated by calculating the Euclidean distances between point coordinates (sampled at *Fs* = 2Hz) for each individual gameplay-trajectory recorded at the first attempt of the relevant game level. It should be noted here that, during the game, players were allowed to repeat levels as many times as they wished and, for that reason, the scope of the analysis was limited to the first attempts only.

Following Coutrot et al. [19], the distances of trajectories in relevant wayfinding tasks were then corrected by dividing them by sum of mean Euclidean distances travelled at Levels 1 and 2 across up to first three attempts (corrected distances). As some participants played Levels 1 and 2 only once but some others approached them multiple times, we considered only up to the first three attempts to control for any undesired outcomes or technical issues encountered in these early tutorial levels. The obtained corrected distances for each wayfinding level were additionally scaled (*i.e. z*-score standardised) to allow unbiased comparison between levels of distinct spatial layouts.

We then calculated the average (i.e. arithmetic mean) distance travelled by each individual player across six wayfinding tasks used in this analysis by simply summing up individual corrected and standardised distances obtained for each level and then dividing it by a number of levels each player completed. Finally, as the estimated metric was in fact a mean distance travelled from the origin to the destination (*i.e.* the smaller the value, the better the performance), we reversed its sign for easier interpretation (*i.e.* the larger the value, the better the performance). It represented a single and unbiased measurement of one’s wayfinding performance across multiple wayfinding tasks.

Wayfinding performance for countries and cultural clusters was estimated with conditional modes by extracting the intercepts for 46 countries in the sample and 11 cultural clusters from linear mixed models with fixed effects for age and gender and random effect for country (or cluster, respectively): Wayfinding Performance ~ Age + Gender + (1 / Country) and Wayfinding Performance ~ Age + Gender + (1 / Cluster). We used lme4 [59] and lmerTest [60] packages for the R programming language to implement these models.

#### Measuring the gap between self-estimated navigation skills and wayfinding performance

The gap between self-reported navigation skills and wayfinding performance was estimated as the difference between the min-max normalised (range [0, 1]) self-estimated ability for each country or a cultural cluster (i.e. average self-estimates controlled for age and gender, extracted as country/cluster conditional modes from the linear mixed models described earlier) and the min-max normalised (range [0, 1]) wayfinding performance (i.e. average wayfinding performance controlled for age and gender, extracted as country/cluster conditional modes from the linear mixed models defined above). The resulting gap metric was distributed on the scale with range of [−1, 1], where −1 denoted maximum possible underestimation of navigation skills compared to the measured wayfinding performance, value 0 indicated perfectly accurate estimation of navigation skills across country/cluster, whereas value 1 expressed the maximum possible overestimation of wayfinding abilities.

#### Data processing, sampling and controlling for fake demographics

As not every participant played all game levels, the sample sizes used to test specific research questions varied quite considerably depending on which levels were used in the analysis. The final sample size was also restricted by selecting only those players who provided the self-estimate scores of their navigation skills and by narrowing the age range of individuals to between 19 and 70 years old in order to control for fake age demographics as well as previously reported selection bias [19] which was found to translate to an unusual spike in performance in older players. The sample was additionally adjusted by a.) removing outliers (defined as players who scored at least two standard deviations above and below the mean of wayfinding performance metric (see above), and b.) keeping the records of only those individuals who represented countries with at least 500 players. Following these data processing operations, the resulting sample size for testing the hypotheses was 383187 individuals representing 46 countries. Detailed descriptives for each country are presented in Supplementary Information, Appendix A, Table S1.

Also, due to the applied standardisation and outlier removal approaches, the normality assumptions were met for some analyses but not for others, and therefore our methods implemented a range of parametric and non-parametric statistical techniques depending on underlying distributions of wayfinding performance metrics, specific cultural dimensions, global indices and particular nature of research questions. These are indicated in the main text of the manuscript and the supplementary appendices whenever statistical tests are reported.

## Data availability

Access to the full Sea Hero Quest dataset (including all wayfinding trajectories generated by the players) is available upon registration through a dedicated server at https://shqdata.z6.web.core.windows.net/.

Please contact the Lead Contacts directly (H.J.S. and E.M.) to obtain the sample of the full dataset used in this study.

Lead Contacts: Hugo J. Spiers and Ed Manley.

Additionally, researchers interested in using the Sea Hero Quest mobile game application can invite participants to play the game and generate wayfinding research data for non-commercial purposes via the SHQ game portal at https://seaheroquest.alzheimersresearchuk.org/.

Access information to secondary data sources can be obtained from the following URLs:

Hofstede’s cultural dimensions (2016) - https://geerthofstede.com/research-and-vsm/dimension-data-matrix/.

Gallup World Poll 2005-2017 (Life Satisfaction) - https://www.gallup.com/178667/gallup-world-poll-work.aspx.

Gallup Religiosity Index (2009) - https://rationalwiki.org/wiki/Importance_of_religion_by_country.

United Nations Development Programme (Education Index 2018, Gender Development Index 2019, and Gender Inequality Index 2018) - http://hdr.undp.org/en/data.

Global Gender Gap Index 2020 - https://www.weforum.org/reports/gender-gap-2020-report-100-years-pay-equality.

GDP per capita (2019) - https://data.worldbank.org/.

## Code availability

All data pre-processing operations, statistical analyses and figures included in this manuscript were made using the R and Python programming languages. The custom code scripts are available from the Lead Contacts (H.J.S. and E.M.) upon request.

## Ethics declaration

All participants voluntarily downloaded and played the Sea Hero Quest mobile application game. This study was conducted as part of a larger research project which has been approved by the UCL Ethics Research Committee under the project number: CPB/2013/015. The authors declare no competing interests.

## Author Contributions

Conceptualization, S.W., H.J.S, M.Ho., and E.M. Methodology, S.W, H.J.S., and E.M. Investigation, S.W., H.J.S., and E.M. Formal analysis, S.W., H.J.S., and E.M. Resources, H.J.S, M.Ho., R.C.D., J.M.W., and C.H. Data curation, S.W., H.J.S, M.Ho., R.C.D., J.M.W., and C.H. Writing - Original Draft, S.W., H.J.S, and E.M. Writing - Review & Editing, S.W., A.C., M.He., P.F.V., H.J.S., and E.M. with input from all authors.

Visualization, S.W., H.J.S, and E.M. Supervision, H.J.S, and E.M.

## Acknowledgments

This manuscript was written as part of the Ph.D. research conducted by S.W. whose work was funded by the Alan Turing Institute in London. S.W. would like to thank the Alan Turing Institute for academic support and financial contribution during his doctoral research. Furthermore, the authors wish to thank Deutsche Telekom and Alzheimer’s Research UK (ARUK-DT2016-1) for funding the research and analysis based on the Sea Hero Quest mobile game application. Finally, the authors are grateful to Glitchers Limited for the SHQ game production, and to Saatchi & Saatchi London for the project management and creative input.

## References

1. Kozlowski, L.T., and Bryant, K.J. (1977). Sense of direction, spatial orientation, and cognitive maps. J. Exp. Psychol. Hum. Percept. Perform 3(4), 590–598. 10.1037/0096-1523.3.4.590

2. Sholl, M.J., Kenny, R.J., and DellaPorta, K.A. (2006). Allocentric-heading recall and its relation to self-reported sense-of-direction. J. Exp. Psychol. Learn. Mem. Cogn. 32(3), 516–533. 10.1037/0278-7393.32.3.516

3. Epstein, R.A., Higgins, J.S., and Thompson-Schill, S.L. (2005). Learning places from views: variation in scene processing as a function of experience and navigational ability. J. Cogn. Neurosci 17(1), 73–83. 10.1162/0898929052879987

4. Hegarty, M., Burte, H., and Boone, A.P. (2018). Individual differences in large-scale spatial abilities and strategies. In Handbook of behavioral and cognitive geography, D. R. Montello, ed. (Edward Elgar Publishing), pp. 231–246. 10.4337/9781784717544.00022

5. Hegarty, M., Richardson, A.E., Montello, D.R., Lovelace, K., and Subbiah, I. (2002). Development of a self-report measure of environmental spatial ability. Intelligence 30, 425–447. 10.1016/S0160-2896(02)00116-2

6. Mitolo, M., Gardini, S., Caffarra, P., Ronconi, L., Venneri, A., and Pazzaglia, F. (2015). Relationship between spatial ability, visuospatial working memory and self-assessed spatial orientation ability: A study in older adults. Cogn. Process. 16(2), 165–176. 10.1007/s10339-015-0647-3

7. Linn, M.C., and Petersen, A.C. (1985). Emergence and characterization of sex differences in spatial ability: A meta-analysis. Child Dev. 56(6), 1479–1498. 10.2307/1130467

8. Nazareth, A., Huang, X., Voyer, D., and Newcombe, N. (2019). A meta-analysis of sex differences in human navigation skills. Psychon. Bull. Rev. 26(5), 1503–1528. 10.3758/s13423-019-01633-6

9. Reilly, D., and Neumann, D.L. (2013). Gender-role differences in spatial ability: A meta-analytic review. Sex Roles 68(9), 521–535. 10.1007/s11199-013-0269-0

10. Munion, A.K., Stefanucci, J.K., Rovira, E., Squire, P., and Hendricks, M. (2019). Gender differences in spatial navigation: Characterizing wayfinding behaviors. Psychon. Bull. Rev. 26(6), 1933–1940. 10.3758/s13423-019-01659-w

11. Weisberg, S. M., Schinazi, V. R., Newcombe, N. S., Shipley, T. F., and Epstein, R. A. (2014). Variations in cognitive maps: understanding individual differences in navigation. J. Exp. Psychol. Learn. Mem. Cogn. 40(3), 669–682. 10.1037/a0035261

12. Saemi, E., Porter, J.M., Ghotbi-Varzaneh, A., Zarghami, M., and Maleki, F. (2012). Knowledge of results after relatively good trials enhances self-efficacy and motor learning. Psychol. Sport. Exerc. 13(4), 378–382. 10.1016/j.psychsport.2011.12.008

13. Wulf, G., Chiviacowsky, S., and Lewthwaite, R. (2012). Altering mindset can enhance motor learning in older adults. Psychol. Aging 27(1), 14–21. 10.1037/a0025718

14. Nietfeld, J.L., Shores, L.R., and Hoffmann, K.F. (2014). Self-regulation and gender within a game-based learning environment. J. Educ. Psychol. 106(4), 961–973. 10.1037/a0037116

15. Bergey, B.W., Ketelhut, D.J., Liang, S., Natarajan, U., and Karakus, M. (2015). Scientific inquiry self-efficacy and computer game self-efficacy as predictors and outcomes of middle school boys’ and girls’ performance in a science assessment in a virtual environment. J. Sci. Educ. Technol. 24(5), 696–708. 10.1007/s10956-015-9558-4

16. Roick, J., and Ringeisen, T. (2017). Self-efficacy, test anxiety, and academic success: A longitudinal validation. Int. J. Educ. Res. 83, 84–93. 10.1016/j.ijer.2016.12.006

17. Bandura, A. (1997). Self-efficacy: The exercise of control (W H Freeman/Times Books/Henry Holt & Co.).

18. Coutrot, A., Manley, E., Goodroe, S., Gahnstrom, C., Filomena, G., Yesiltepe, D., Dalton, R.C., Wiener, J.M., Hölscher, C., Hornberger, M., and Spiers, H.J. (2022). Entropy of city street networks linked to future spatial navigation ability. Nature 604(7904), 104–110. 10.1038/s41586-022-04486-7

19. Coutrot, A., Silva, R., Manley, E., de Cothi, W., Sami, S., Bohbot, V.D., Wiener, J.M., Hölscher, C., Dalton, R.C., Hornberger, M., and Spiers, H.J. (2018). Global determinants of navigation ability. Curr. Biol. 28(17), 2861–2866. 10.1016/j.cub.2018.06.009

20. Castelli, L., Latini Corazzini, L., and Geminiani, G.C. (2008). Spatial navigation in large-scale virtual environments: Gender differences in survey tasks. Comput. Hum. Behav. 24(4), 1643–1667. 10.1016/j.chb.2007.06.005

21. Coluccia, E., and Louse, G. (2004). Gender differences in spatial orientation: A review. J. Environ. Psychol. 24(3), 329–340. 10.1016/j.jenvp.2004.08.006

22. Taillade, M., N’Kaoua, B., and Sauzéon, H. (2016). Age-related differences and cognitive correlates of self-reported and direct navigation performance: The effect of real and virtual test conditions manipulation. Front. Psychol. 6(2034). 10.3389/fpsyg.2015.02034

23. Yuan, L., Kong, F., Luo, Y., Zeng, S., Lan, J., and You, X. (2019). Gender differences in large-scale and small-scale spatial ability: A systematic review based on behavioral and neuroimaging research. Front. Behav. Neurosci. 13(128). 10.3389/fnbeh.2019.00128

24. Zancada-Menendez, C., Sampedro-Piquero, P., Lopez, L., and McNamara, T.P. (2016). Age and gender differences in spatial perspective taking. Aging Clin. Exp. Res. 28, 289–296. 10.1007/s40520-015-0399-z

25. Weisberg, S.M., Schinazi, V.R., Newcombe, N.S., Shipley, T.F., and Epstein, R.A. (2014). Variations in cognitive maps: Understanding individual differences in navigation. J. Exp. Psychol. Learn. Mem. Cogn. 40(3), 669–682. 10.1037/a0035261

26. Boone, A.P., Gong, X., and Hegarty, M. (2018). Sex differences in navigation strategy and efficiency. Mem. Cogn. 46, 909–922. 10.3758/s13421-018-0811-y

27. Hegarty, M., He, C., Boone, A.P., Yu, S., Jacobs, E.G., and Chrastil, E.R. (2022). Understanding differences in wayfinding strategies. Top. Cogn. Sci. 10.1111/tops.12592

28. Yu, S., Boone, A.P., He, C., Davis, R.C., Hegarty, M., Chrastil, E.R., and Jacobs, E.G. (2021). Age-related changes in spatial navigation are evident by midlife and differ by sex. Psychol. Sci. 32(5), 692–704. 10.1177/0956797620979185

29. Taillade, M., Sauzéon, H., Dejos, M., Pala, P.A., Larrue, F., Wallet, G., Gross, C., and N’Kaoua, B. (2013). Executive and memory correlates of age-related differences in wayfinding performances using a virtual reality application. Neuropsychol. Dev. Cogn. B Aging Neuropsychol. Cogn. 20(3), 298–319. 10.1080/13825585.2012.706247

30. Van der Ham, I.J.M., Van der Kuil, M.N.A, and Claessen, M.H.G. (2020). Quality of self-reported cognition: Effects of age and gender on spatial navigation self-reports. Aging Ment. Health 25(5), 873–878. 10.1080/13607863.2020.1742658

31. van der Ham, I.J.M., and Koutzmpi, V. (2022). Stereotypes and self-reports about spatial cognition: Impact of gender and age. Curr. Psychol. 10.1007/s12144-022-03827-z

32. Caffò, A.O., Lopez, A., Spano, G., Serino, S., Cipresso. P., Stasolla, F., Savino, M., Lancioni, G.E., Riva, G., and Bosco, A. (2018). Spatial reorientation decline in aging: The combination of geometry and landmarks. Aging Ment. Health 22, 1372–1383. 10.1080/13607863.2017.1354973

33. De Beni, R., Pazzaglia, F., and Gardini, S. (2006). The role of mental rotation and age in spatial perspective-taking tasks: When age does not impair perspective-taking performance. Appl. Cogn. Psychol. 20(6), 807–821. 10.1002/acp.1229

34. Hegarty, M., Montello, D.R., Richardson, A.E., Ishikawa, T., and Lovelace, K. (2006). Spatial abilities at different scales: Individual differences in aptitude-test performance and spatial-layout learning. Intelligence, 34(2), 151–176. 10.1016/j.intell.2005.09.005

35. Nori, R., and Piccardi, L. (2015). I believe I’m good at orienting myself… But is that true? Cogn. Process. 16, 301–307. 10.1007/s10339-015-0655-3

36. Cross, C.P., Brown, G.R., Morgan, T.J.H., and Laland, K.N. (2017). Sex differences in confidence influence patterns of conformity. Br. J. Psychol. 108(4), 655–667. 10.1111/bjop.12232

37. Estes, Z., and Felker, S. (2012). Confidence mediates the sex difference in mental rotation performance. Arch. Sex. Behav. 41(3), 557–570. 10.1007/s10508-011-9875-5

38. Lawton, C.A., Charleston, S.I., and Zieles, A.S. (1996). Individual- and gender-related differences in indoor wayfinding. Environ. Behav. 28(2), 204–219. 10.1177/0013916596282003

39. Picucci, L., Caffò, A.O., and Bosco, A. (2011). Besides navigation accuracy: Gender differences in strategy selection and level of spatial confidence. J. Environ. Psychol. 31(4), 430–438. 10.1016/j.jenvp.2011.01.005

40. Vander Heyden, K.M., van Atteveldt, N.M., Huizinga, M., and Jolles, J. (2016). Implicit and explicit gender beliefs in spatial ability: Stronger stereotyping in boys than girls. Front. Psychol. 7(1114). 10.3389/fpsyg.2016.01114

41. Verde, P., Piccardi, L., Bianchini, F., Trivelloni, P., Guariglia, C., and Tomao, E. (2013). Gender effects on mental rotation in pilots vs. nonpilots. Aviat. Space Environ. Med. 84(7), 726–729. 10.3357/asem.3466.2013

42. Verde, P., Piccardi, L., Bianchini, F., Guariglia, C., Carrozzo, P., Morgagni, F., Boccia, M., Di Fiore, G., and Tomao, E. (2015). Gender differences in navigational memory: pilots vs. nonpilots. Aerosp. Med. Hum. Perform. 86(2), 103–111. 10.3357/AMHP.4024.2015

43. Diekman, A.B., and Eagly, A.H. (2000). Stereotypes as dynamic constructs: Women and men of the past, present, and future. Pers. Soc. Psychol. Bull. 26(10), 1171–1188. 10.1177/0146167200262001

44. Yang, Y., and Merrill, E.C. (2017). Cognitive and personality characteristics of masculinity and femininity predict wayfinding competence and strategies of men and women. Sex Roles, 76(11-12), 747–758. 10.1007/s11199-016-0626-x

45. Oettingen, G. (1995). Cross-cultural perspectives on self-efficacy. In Self-Efficacy in Changing Societies, A. Bandura, ed. (Cambridge University Press), pp. 149–176. 10.1017/CBO9780511527692.007

46. Hofstede, G. (2001). Culture’s Consequences: Comparing Values, Behaviors, Institutions, and Organizations Across Nations (Sage).

47. Hofstede, G., Hofstede, G.J., and Minkov, M. (2010). Cultures and organizations: Software of the mind; intercultural cooperation and its importance for survival (McGraw-Hill).

48. Klassen, R.M. (2004). Optimism and realism: A review of self-efficacy from a cross-cultural perspective. Int. J. Psychol. 39(3), 205–230. 10.1080/00207590344000330

49. Luszczynska, A., Gutiérrez-Doña, B., and Schwarzer, R. (2005). General self-efficacy in various domains of human functioning: Evidence from five countries. Int. J. Psychol. 40(2), 80–89. 10.1080/00207590444000041

50. Camgoz, S.M., Tektas, O.O., and Metin, I. (2008). Academic attributional style, self-efficacy and gender: A cross-cultural comparison. Soc. Behav. Pers. 36(1), 97–114. 10.2224/sbp.2008.36.1.97

51. Lee, J. (2009). Self-Constructs and Anxiety Across Cultures. ETS Res. Rep. Ser. 2009(1), i–35. 10.1002/j.2333-8504.2009.tb02169.x

52. Chiu, M.M., and Klassen, R.M. (2010). Relations of mathematics self-concept and its calibration with mathematics achievement: Cultural differences among fifteen-year-olds in 34 countries. Learn. Instr. 20(1), 2–17. 10.1016/j.learninstruc.2008.11.002

53. Jurecska, D.E.S., Lee, C.E., Chang, K.B.T., and Sequeira, E. (2011). I am smart, therefore I can: Examining the relationship between IQ and self-efficacy across cultures. J. Adolesc. Health 23(3), 209–216. 10.1515/ijamh.2011.046

54. Bonneville-Roussy, A., Bouffard, T., Palikara, O., and Vezeau, C. (2019). The role of cultural values in teacher and student self-efficacy: Evidence from 16 nations. Contemp. Educ. Psychol. 59, 101798. 10.1016/j.cedpsych.2019.101798

55. Zusho, A., and Clayton, K.E. (2015). Cross-Cultural Study of Education. In International Encyclopedia of the Social & Behavioral Sciences, J. D. Wright, ed. (Elsevier), pp. 327–333.

56. Ronen, S., and Shenkar, O. (2013). Mapping world cultures: Cluster formation, sources and implications. J. Int. Bus. Stud. 44, 867–897. 10.1057/jibs.2013.42

57. Coutrot, A., Schmidt, S., Coutrot, L., Pittman, J., Hong, L., Wiener, J.M., Hölscher, C., Dalton, R.C., Hornberger, M., and Spiers, H.J. (2019). Virtual navigation tested on a mobile app is predictive of real-world wayfinding navigation performance. PLoS One 14(3), e0213272. 10.1371/journal.pone.0213272

58. House, R.J., Hanges, P.J., Javidan, M., Dorfman, P.W., and Gupta, V. (2004). Culture, Leadership, and Organizations: The GLOBE Study of 62 Societies (SAGE Publications).

59. Bates, D., Mächler, M., Bolker, B., and Walker, S. (2015). Fitting Linear Mixed-Effects Models Using lme4. J. Stat. Softw. 67(1), 1–48. 10.18637/jss.v067.i01

60. Kuznetsova, A., Brockhoff, P.B., and Christensen, R.H.B. (2017). lmerTest Package: Tests in Linear Mixed Effects Models. J. Stat. Softw. 82(13), 1–26. 10.18637/jss.v082.i13

61. Majid, A., Bowerman, M., Kita, S., Haun, D. B., and Levinson, S. C. (2004). Can language restructure cognition? The case for space. Trends Cogn. Sci. 8(3), 108–114. 10.1016/j.tics.2004.01.003

62. Barhorst-Cates, E. M., Meneghetti, C., Zhao, Y., Pazzaglia, F., and Creem-Regehr, S. H. (2021). Effects of home environment structure on navigation preference and performance: A comparison in Veneto, Italy and Utah, USA. J. Environ. Psychol. 74, 101580. 10.1016/j.jenvp.2021.101580

63. Davis, H. E., Stack, J., and Cashdan, E. (2021). Cultural change reduces gender differences in mobility and spatial ability among seminomadic pastoralist-forager children in northern Namibia. Hum. Nat. 32(1), 178–206. 10.1007/s12110-021-09388-7

64. Davis, H. E., Gurven, M., and Cashdan, E. (2022). Navigational Experience and the Preservation of Spatial Abilities into Old Age Among a Tropical Forager-Farmer Population. Top. Cogn. Sci. 10.1111/tops.12602

65. Schug, M. G., Barhorst-Cates, E., Stefanucci, J., Creem-Regehr, S., Olsen, A. P., and Cashdan, E. (2022). Childhood Experience Reduces Gender Differences in Spatial Abilities: A Cross-Cultural Study. Cogn. Sci. 46(2), e13096. 10.1111/cogs.13096

