## Supplementary Information for "Cultural determinants of the gap between self-estimated navigation ability and wayfinding performance: evidence from 46 countries"

### *Appendix A - Tables and Figures*

*Table S1. Descriptive statistics by country and gender for the sample of 383,187 players representing 46 countries used in this study.*

| | | | | | Females | | | Males | | |
|---|---|---|---|---|---|---|---|---|---|---|
| | Country | N | Mean age | SD age | N | Mean age | SD age | N | Mean age | SD age |
| 1 | Argentina | 1774 | 33.87 | 12.61 | 671 | 33.92 | 13.44 | 1103 | 33.83 | 12.08 |
| 2 | Australia | 7952 | 39.29 | 14.36 | 4031 | 39.35 | 14.98 | 3921 | 39.22 | 13.69 |
| 3 | Austria | 1974 | 36.66 | 13.51 | 809 | 34.98 | 14.19 | 1165 | 37.83 | 12.89 |
| 4 | Belgium | 1941 | 38.6 | 14.56 | 778 | 39.24 | 15.13 | 1163 | 38.17 | 14.16 |
| 5 | Brazil | 6105 | 32.66 | 12.4 | 2676 | 32.28 | 13.27 | 3429 | 32.96 | 11.67 |
| 6 | Canada | 10851 | 40.19 | 14.77 | 5187 | 39.99 | 15.45 | 5664 | 40.36 | 14.11 |
| 7 | Chile | 1606 | 33.61 | 12.68 | 640 | 31.88 | 12.93 | 966 | 34.76 | 12.38 |
| 8 | China | 5189 | 27.68 | 7.53 | 2580 | 26.94 | 7.23 | 2609 | 28.42 | 7.74 |
| 9 | Colombia | 1457 | 31.04 | 11 | 551 | 30.63 | 11.57 | 906 | 31.3 | 10.63 |
| 10 | Costa Rica | 596 | 34.16 | 12.24 | 256 | 33.83 | 13.18 | 340 | 34.41 | 11.48 |
| 11 | Czech Republic | 17023 | 29.9 | 10.6 | 7958 | 29.33 | 10.99 | 9065 | 30.4 | 10.22 |
| 12 | Denmark | 1714 | 37.08 | 13.12 | 585 | 35.88 | 13.8 | 1129 | 37.7 | 12.71 |
| 13 | Finland | 704 | 35.84 | 12.25 | 269 | 35.06 | 12.8 | 435 | 36.33 | 11.88 |
| 14 | France | 7943 | 36.74 | 13.94 | 3285 | 36.8 | 14.94 | 4658 | 36.69 | 13.19 |
| 15 | Germany | 34495 | 40.71 | 13.51 | 15632 | 40.15 | 13.9 | 18863 | 41.18 | 13.15 |
| 16 | Greece | 23745 | 32.84 | 10.59 | 9355 | 31.83 | 10.73 | 14390 | 33.49 | 10.44 |
| 17 | Hong Kong | 989 | 34.12 | 11.97 | 379 | 31.41 | 11.48 | 610 | 35.8 | 11.96 |
| 18 | Hungary | 10728 | 31.21 | 11.22 | 4748 | 30.52 | 11.41 | 5980 | 31.76 | 11.03 |
| 19 | India | 3341 | 26.73 | 8.14 | 496 | 29.13 | 10.65 | 2845 | 26.31 | 7.54 |
| 20 | Indonesia | 1095 | 27.3 | 8.47 | 330 | 26.23 | 7.54 | 765 | 27.76 | 8.8 |
| 21 | Ireland | 1408 | 39.43 | 13.13 | 546 | 39.01 | 14.36 | 862 | 39.7 | 12.29 |
| 22 | Islamic Republic of Iran | 1071 | 31.99 | 8.97 | 281 | 31.04 | 9.64 | 790 | 32.32 | 8.7 |
| 23 | Italy | 11780 | 35.39 | 13.56 | 4573 | 34.2 | 13.75 | 7207 | 36.14 | 13.38 |
| 24 | Malaysia | 1161 | 29.04 | 10.48 | 444 | 28.14 | 10.16 | 717 | 29.6 | 10.64 |
| 25 | Mexico | 3923 | 29.51 | 10.73 | 1492 | 28.64 | 10.74 | 2431 | 30.04 | 10.69 |
| 26 | Netherlands | 22877 | 35.97 | 13.43 | 12490 | 35.9 | 13.75 | 10387 | 36.05 | 13.04 |
| 27 | New Zealand | 1700 | 40.07 | 14.76 | 939 | 40.23 | 15.44 | 761 | 39.88 | 13.88 |
| 28 | Norway | 1140 | 35.25 | 12.33 | 483 | 32.79 | 12.13 | 657 | 37.05 | 12.17 |
| 29 | Philippines | 1370 | 27.86 | 9.88 | 673 | 26.8 | 9.19 | 697 | 28.88 | 10.4 |
| 30 | Poland | 9209 | 29.34 | 9.73 | 4496 | 28.52 | 9.89 | 4713 | 30.13 | 9.51 |
| 31 | Portugal | 2077 | 34.65 | 11.02 | 850 | 32.89 | 11.14 | 1227 | 35.86 | 10.78 |
| 32 | Romania | 3208 | 29.82 | 9.11 | 1144 | 28.1 | 8.91 | 2064 | 30.78 | 9.08 |
| 33 | Russian Federation | 2761 | 27.58 | 7.98 | 948 | 26.18 | 8.24 | 1813 | 28.32 | 7.74 |
| 34 | Singapore | 1338 | 32.48 | 11.49 | 529 | 31.22 | 11.18 | 809 | 33.3 | 11.62 |
| 35 | Slovakia | 5096 | 29.31 | 10.14 | 2297 | 28.7 | 10.28 | 2799 | 29.81 | 10 |
| 36 | South Africa | 1126 | 37.92 | 13.56 | 455 | 38.58 | 14.24 | 671 | 37.47 | 13.07 |
| 37 | Spain | 6797 | 36.99 | 13 | 2661 | 36.43 | 13.46 | 4136 | 37.36 | 12.68 |
| 38 | Sweden | 1793 | 35.85 | 12.72 | 667 | 34.55 | 12.88 | 1126 | 36.62 | 12.56 |
| 39 | Switzerland | 3136 | 41.82 | 14.1 | 1330 | 40.51 | 14.78 | 1806 | 42.78 | 13.51 |
| 40 | Taiwan | 1511 | 32.34 | 10.72 | 634 | 31.71 | 11.45 | 877 | 32.8 | 10.15 |
| 41 | Thailand | 871 | 30.88 | 11.67 | 322 | 28.83 | 10.62 | 549 | 32.08 | 12.1 |
| 42 | Turkey | 1835 | 31.19 | 9.75 | 602 | 29.35 | 9.61 | 1233 | 32.09 | 9.69 |
| 43 | Ukraine | 647 | 27.59 | 8.29 | 181 | 26.99 | 8.53 | 466 | 27.83 | 8.2 |
| 44 | United Arab Emirates | 685 | 35.68 | 11.1 | 197 | 33.92 | 11.6 | 488 | 36.38 | 10.82 |
| 45 | United Kingdom | 66196 | 42.94 | 14.11 | 28727 | 42.68 | 14.73 | 37469 | 43.14 | 13.61 |
| 46 | United States | 87249 | 38.99 | 14.55 | 43792 | 39.77 | 15.15 | 43457 | 38.21 | 13.89 |

*Table S2. Left - odds ratios and t-values (along with p-values) for the multinomial logistic regression model with the self-estimated navigation skills (categorical) as the dependent variable and age band (categorical) as well as gender (categorical) as the independent variables. Right - probabilities of self-estimated navigation skills by gender and age band obtained from the logistic regression model.*

| Coefficients | Odds Ratios | t value |
|---|---|---|
| **Predictors:** | | |
| Age band (reference level: *19-29*) | | |
| *30-39* | 1.22 | 22.89*** |
| *40-49* | 1.30 | 29.24*** |
| *50-59* | 1.30 | 26.30*** |
| *60-70* | 1.12 | 8.97*** |
| Gender (reference level: *female*) | | |
| *male* | 2.61 | 145.16*** |
| **Intercepts:** | | |
| *very bad \| bad* | 0.03 | -264.99*** |
| *bad \| good* | 0.24 | -209.45*** |
| *good \| very good* | 4.03 | 208.41*** |
| $R^2$ Nagelkerke | 0.070 | |

*$p < 0.05$; **$p < 0.01$; ***$p < 0.001$.

| | | | Probabilities of self-estimated navigation skills | | | |
|---|---|---|---|---|---|---|
| | Gender | Age band | very bad | bad | good | very good |
| 1 | female | 19-29 | 0.0291 | 0.1658 | 0.6062 | 0.1989 |
| 2 | male | 19-29 | 0.0113 | 0.0736 | 0.522 | 0.3931 |
| 3 | female | 30-39 | 0.024 | 0.142 | 0.6021 | 0.2318 |
| 4 | male | 30-39 | 0.0094 | 0.0616 | 0.4886 | 0.4405 |
| 5 | female | 40-49 | 0.0224 | 0.1341 | 0.5988 | 0.2447 |
| 6 | male | 40-49 | 0.0087 | 0.0577 | 0.4756 | 0.458 |
| 7 | female | 50-59 | 0.0226 | 0.1346 | 0.5991 | 0.2437 |
| 8 | male | 50-59 | 0.0088 | 0.058 | 0.4766 | 0.4567 |
| 9 | female | 60-70 | 0.0261 | 0.1522 | 0.6048 | 0.2168 |
| 10 | male | 60-70 | 0.0102 | 0.0666 | 0.5038 | 0.4194 |

*Table S3. Hierarchical regression analysis of predictors of wayfinding performance. Model 1 (without the self-estimates of navigation skills) is significantly weaker in terms of explained variance than Model 2 (with the self-estimates of navigation skills). Model 2 shows that the self-estimated navigation skills variable is a significant predictor of wayfinding performance measured across six wayfinding tasks (SHQ game levels: 3, 6, 7, 8, 11, and 12).*

| Predictor variables | Model 1 | | Model 2 | |
|---|---|---|---|---|
| | standardized ß coefficient | *t* value | standardized ß coefficient | *t* value |
| Intercept | -0.399 | 40.544*** | -0.567 | 38.388*** |
| Age | -0.352 | 232.692*** | -0.355 | 234.788*** |
| Gender (reference level: *female*) | | | | |
| *male* | 0.462 | -153.757*** | 0.433 | -140.304*** |
| Education (reference level: *no formal*) | | | | |
| *high school* | 0.105 | -10.976*** | 0.099 | -10.409*** |
| *college* | 0.187 | -19.658*** | 0.179 | -18.928*** |
| *university* | 0.207 | -22.211*** | 0.199 | -21.339*** |
| Travel time (reference level: *up to 30 mins*) | | | | |
| *30 mins to 1 hour* | -0.008 | 2.276* | -0.012 | 3.555*** |
| *1 hour+* | -0.030 | 7.708*** | -0.042 | 10.949*** |
| Home environment (reference level: *rural*) | | | | |
| *mixed* | 0.028 | -6.953*** | 0.031 | -7.749*** |
| *city* | -0.090 | 20.230*** | -0.086 | 19.394*** |
| Navigation skills (reference level: *very bad*) | | | | |
| *bad* | - | - | 0.078 | -6.331*** |
| *good* | - | - | 0.178 | -15.364*** |
| *very good* | - | - | 0.263 | -22.440*** |
| $R^2$ | 0.1789 | | 0.1824 | |
| $R^2$ change | - | | 0.0035 | |

*$p < 0.05$; **$p < 0.01$; ***$p < 0.001$.

*Table S4. ANOVA tables of terms in the linear regression models without self-estimates of navigation skills (Left), with self-estimates of navigation skills (Right), and for the F-test of differences between both models (Bottom).*

| | Sum Sq | Df | F value | Pr(>F) |
|---|---|---|---|---|
| Age | 44332.08 | 1 | 54145.661 | 0 |
| Gender | 19356.316 | 1 | 23641.131 | 0 |
| Education | 944.064 | 3 | 384.349 | 0 |
| Travel time | 50.244 | 2 | 30.683 | 0 |
| Home environment | 974.059 | 2 | 594.841 | 0 |
| Residuals | 303401.465 | 370564 | | |

| | Sum Sq | Df | F value | Pr(>F) |
|---|---|---|---|---|
| Age | 44939.321 | 1 | 55125.378 | 0 |
| Gender | 16047.676 | 1 | 19685.082 | 0 |
| Education | 883.151 | 3 | 361.109 | 0 |
| Travel time | 99.894 | 2 | 61.268 | 0 |
| Home environment | 957.642 | 2 | 587.352 | 0 |
| Navigation skills | 1312.654 | 3 | 536.728 | 0 |
| Residuals | 302088.811 | 370561 | | |

| | Res.Df | RSS | Df | Sum of Sq | F | Pr(>F) |
|---|---|---|---|---|---|---|
| 1 | 370564 | 303401.465 | | | | |
| 2 | 370561 | 302088.811 | 3 | 1312.654 | 536.728 | 0 |

*Table S5. Proportions of self-reported navigation skills in the sample by country.*

| | | Proportions of self-estimated navigation skills | | | |
|---|---|---|---|---|---|
| | Country | very bad | bad | good | very good |
| 1 | Argentina | 1.7% | 8.4% | 49.3% | 40.6% |
| 2 | Australia | 2.1% | 11.7% | 56.4% | 29.8% |
| 3 | Austria | 0.6% | 3.8% | 37.9% | 57.7% |
| 4 | Belgium | 2.4% | 11.9% | 58.9% | 26.8% |
| 5 | Brazil | 2.3% | 10.9% | 53.4% | 33.3% |
| 6 | Canada | 2.0% | 11.5% | 52.9% | 33.7% |
| 7 | Chile | 1.5% | 9.3% | 52.5% | 36.7% |
| 8 | China | 3.9% | 16.4% | 57.3% | 22.4% |
| 9 | Colombia | 1.3% | 10.1% | 54.2% | 34.4% |
| 10 | Costa Rica | 2.2% | 9.2% | 49.0% | 39.6% |
| 11 | Czech Republic | 0.8% | 7.6% | 59.3% | 32.3% |
| 12 | Denmark | 2.4% | 11.0% | 53.2% | 33.4% |
| 13 | Finland | 2% | 13% | 60% | 25% |
| 14 | France | 2% | 14% | 59% | 24% |
| 15 | Germany | 0.3% | 3.4% | 42.8% | 53.4% |
| 16 | Greece | 1.7% | 10.3% | 51.6% | 36.4% |
| 17 | Hong Kong | 2% | 15% | 55% | 27% |
| 18 | Hungary | 1.9% | 10.5% | 59.0% | 28.6% |
| 19 | India | 1.9% | 10.9% | 58.7% | 28.6% |
| 20 | Indonesia | 1% | 12% | 60% | 27% |
| 21 | Iran | 2% | 14% | 57% | 26% |
| 22 | Ireland | 1.7% | 9.8% | 56.5% | 32.0% |
| 23 | Italy | 2% | 12% | 55% | 31% |
| 24 | Malaysia | 2.4% | 15.5% | 61.0% | 21.1% |
| 25 | Mexico | 1.3% | 9.6% | 55.8% | 33.3% |
| 26 | Netherlands | 2.0% | 14.5% | 62.7% | 20.8% |
| 27 | New Zealand | 1.9% | 10.7% | 58.9% | 28.5% |
| 28 | Norway | 2% | 12% | 60% | 26% |
| 29 | Philippines | 1.2% | 10.8% | 63.0% | 25.0% |
| 30 | Poland | 1.1% | 9.4% | 56.8% | 32.7% |
| 31 | Portugal | 2.3% | 11.6% | 59.1% | 27.1% |
| 32 | Romania | 1.0% | 6.9% | 59.8% | 32.3% |
| 33 | Russian Federation | 1.4% | 10.5% | 54.2% | 33.9% |
| 34 | Singapore | 3% | 14% | 56% | 26% |
| 35 | Slovakia | 1.1% | 6.2% | 58.0% | 34.8% |
| 36 | South Africa | 2.3% | 10.9% | 51.6% | 35.2% |
| 37 | Spain | 2% | 12% | 55% | 30% |
| 38 | Sweden | 1.6% | 11.6% | 56.3% | 30.4% |
| 39 | Switzerland | 1.08% | 6.82% | 46.33% | 45.76% |
| 40 | Taiwan | 2.2% | 16.4% | 59.8% | 21.6% |
| 41 | Thailand | 2.4% | 12.3% | 60.3% | 25.0% |
| 42 | Turkey | 2% | 13% | 53% | 33% |
| 43 | Ukraine | 1% | 12% | 56% | 30% |
| 44 | United Arab Emirates | 2.6% | 9.9% | 48.0% | 39.4% |
| 45 | United Kingdom | 1.9% | 11.3% | 56.9% | 29.8% |
| 46 | United States | 2.0% | 10.5% | 53.0% | 34.5% |

*Table S6. Sample size and proportions of self-reported navigation skills for each cultural cluster.*

| | Cultural cluster | N | Proportions of self-estimated navigation skills | | | |
|---|---|---|---|---|---|---|
| | | | **very bad** | **bad** | **good** | **very good** |
| 1 | Anglo | 175356 | 2.0% | 10.9% | 54.7% | 32.4% |
| 2 | Eastern Europe | 48672 | 1.2% | 8.6% | 58.3% | 31.9% |
| 3 | Germanic | 39605 | 0.4% | 3.7% | 42.8% | 53.0% |
| 4 | Latin Europe | 30538 | 2% | 13% | 57% | 29% |
| 5 | Nordic | 28228 | 2.0% | 14.0% | 61.5% | 22.5% |
| 6 | Near East | 25580 | 1.7% | 10.4% | 51.7% | 36.2% |
| 7 | Latin America | 15461 | 1.8% | 10.0% | 53.4% | 34.9% |
| 8 | Confucian Asia | 9027 | 3.3% | 16.0% | 57.3% | 23.4% |
| 9 | Far East | 8909 | 2% | 12% | 60% | 26% |
| 10 | African | 1126 | 2.3% | 10.9% | 51.6% | 35.2% |
| 11 | Arabic | 685 | 2.6% | 9.9% | 48.0% | 39.4% |

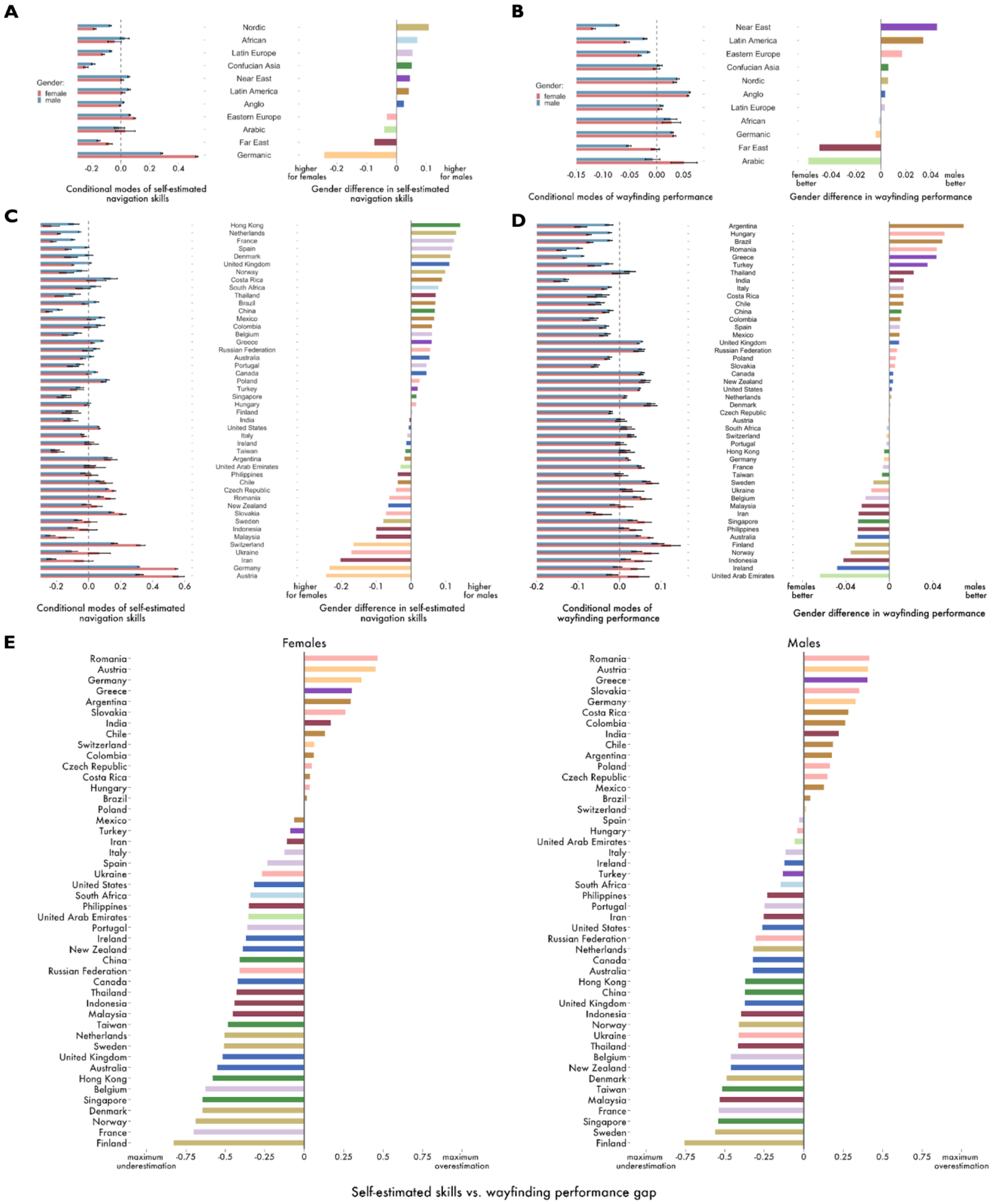


*Figure S1. Self-estimated navigation skills, wayfinding performance and the gap between self-reported skills and actual performance by cultural cluster and country for females and males. A) Conditional modes self-estimated navigation skills by cultural cluster and gender. B) Conditional modes of wayfinding performance by cultural cluster and gender. C) Conditional modes self-estimated navigation skills by country and gender. D) Conditional modes of wayfinding performance by country and gender. E) The gap between self-estimated navigation skills and wayfinding performance by country for females and males.*

*Appendix B - Additional Results*

**The relationships between the self-estimated navigation skills vs. wayfinding performance gap for each country and Hofstede's cultural dimensions as well as global indices.**

**Self-estimated navigation skills vs. wayfinding performance gap by country:** calculated as the difference between min-max normalised navigation skills (estimated as conditional modes obtained from a linear mixed model with self-reported ability as the dependent variable, controlled for age and gender, with varying intercepts for countries) and min-max normalised wayfinding performance (estimated as conditional modes obtained from a linear mixed model with wayfinding performance i.e. reversed MSCD metric as the dependent variable controlled for age and gender, with varying intercepts for countries); range of [-1, 1], where:

- values close to -1 denote 'high underestimation',
- values around 0 mean 'accurate estimation'
- and values closer to 1 translate to 'high overestimation'.

As the gap metric was not normally distributed and most of the global indices and Hofstede's cultural dimensions were also not normally distributed, we estimated the significance of all relationships reported in this appendix with the Spearman's *rho* test.

Correlations between the Self-Estimated Navigation Skills vs. Performance Gap and Hofstede's cultural dimensions:

- Power Distance: Spearman's *rho* = 0.11, p = .491
- Individualism: Spearman's *rho* = -0.18, *p* = .242
- **Masculinity: Spearman's *rho* = 0.43, p = .004****
- **Uncertainty Avoidance: Spearman's *rho* = 0.47, p = .001****
- Long-Term Orientation: Spearman's *rho* = -0.12, *p* = .448
- Indulgence: Spearman's *rho* = -0.18, *p* = .28

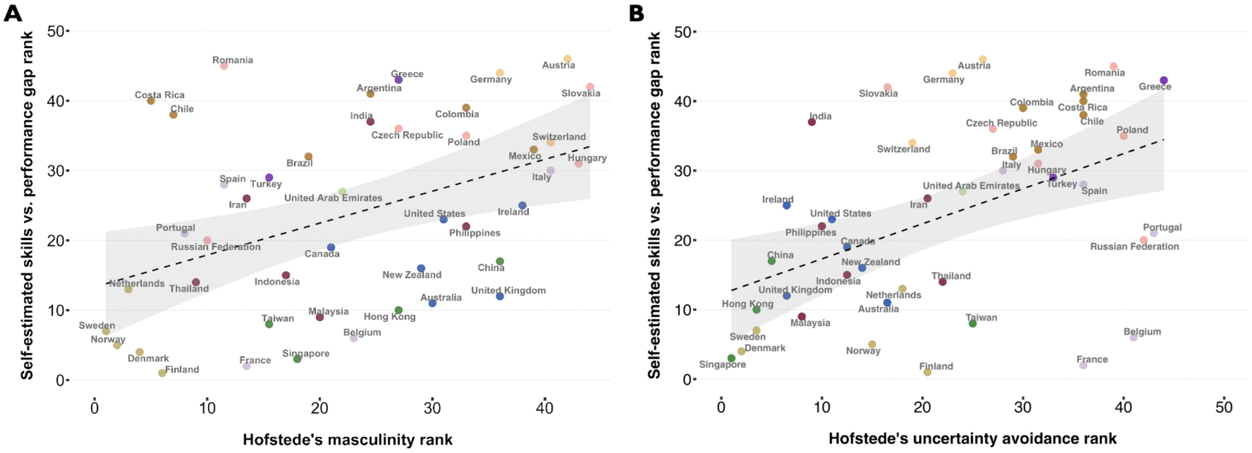


*Figure S2. The Spearman's relationships between the ranks of the self-estimated navigation skills vs. performance gap and the ranks of Hofstede's dimensions of masculinity (A) and uncertainty avoidance (B).*

The above six Hofstede's dimensions were further used in the linear regression model as independent variables of the self-estimated skills vs. performance gap (Table S7).

*Table S7. The effects of Hofstede's cultural dimensions on the self-estimate-performance gap (standardised coefficients and their standard errors with asterisks denoting the level of significance - p-values Bonferroni adjusted). The model $R^2$=50.8% and adjusted $R^2$=42.6%.*

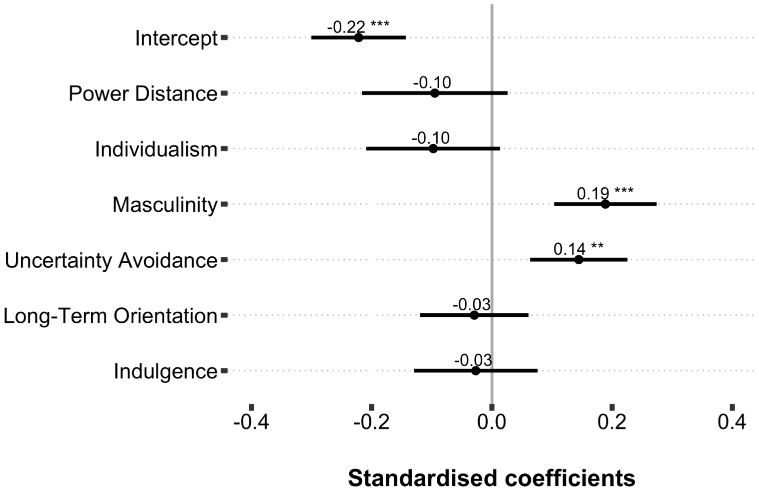


For completeness, we report other significant and non-significant correlations of the gap between the self-estimated navigation skills and wayfinding performance with the remaining global indices used in our study:

Correlation between the Self-Estimated Navigation Skills vs. Performance Gap and the Gender Inequality Index 2018:

- **Gender Inequality Index 2018: Spearman's *rho* = 0.40, *p* = .007****

Correlations between the Self-Estimated Navigation Skills vs. Performance Gap and the Global Gender Gap Report 2020:

- Global Gender Gap Index (overall): Spearman's *rho* = -0.23, *p* = .127
- **Economic Participation & Opportunity: Spearman's *rho* = -0.47, *p* = .001****
- Educational Attainment: Spearman's *rho* = -0.13, *p* = .399
- **Health & Survival: Spearman's *rho* = 0.37, *p* = .01***
- Political Empowerment: Spearman's *rho* = -0.06, *p* = .723

Correlation between the Self-Estimated Navigation Skills vs. Performance Gap and the GDP per capita (2019, World Bank):

- **GDP per capita (2019): Spearman's *rho* = -0.31, *p* = 0.044***

*$p < 0.05$; **$p < 0.01$; ***$p < 0.001$.
